# NaPi- II b as a potential diagnostic and prognostic biomarker in ovarian cancers

**DOI:** 10.1101/394759

**Authors:** Shoufeng Zhao, Zhipeng Wang

## Abstract

Ovarian cancer (OC) is commonly diagnosed at an advanced stage due to a lack of effective biomarkers and specificity required for accurate clinical diagnosis. The purpose of this study was to estimate the diagnosis and prognosis of the NaPi- II b in ovarian cancer. Herein, by performing data mining using the databases of Oncomine and Cancer Cell Line Encyclopedia (CCLE), we are for the first time to report that the expression level of NaPi- II b transcripts in a variety of tumor types compared with the normal controls. Based on Kaplan-Meier plotter, we investigated the prognostic values of NaPi- II b specifically high expressed in OC patients. The results of the Oncomine analysis showed that relative expression of NaPi- II b was distinctly high in OC tissues vs. normal tissues. CCLE analysis indicated that the expression of NaPi- II b in OC cell lines expressed the highest level in all cancer lines. In overall survival (OR) analysis, NaPi- II b mRNA high expressions were correlated to worse OR in OC patients. These results indicate that NaPi- II b may be a novel potential biomarker for determining the diagnosis and predicting the prognosis of OC.

---

Globally, ovarian cancer (OC) remains among the ten top cancers and the most lethal gynecological cancer with dismal prognosis. In 2018, it was estimated that there will be approximately 22,240 new cases and 14,070 deaths from ovarian cancer in the United States (TORRE *et al.* 2018). OC incidence and mortality in China in 2015 are 22,240 and 14,070 respectively (CHEN *et al.* 2016). Owing to a lack of effective biomarkers, OC is usually detected in advanced stages (KOBAYASHI *et al.* 2015). The 5-year survival rate for OC detected in stage I reaches 90% and drops below 30% for stages III-IV (WOLMAN 2014). So, high sensitivity, specificity and good selectivity are need for detection of OC especially for early stage. In the past decades, several biomarkers for OC diagnosis have been studied, such as miR-193a-5p, HE4, CA125, CA19-9 and CEA (GUO *et al.* 2017; REN *et al.* 2018). However, it is not sufficiently specific and diagnostic sensitive to be useful as a screening test (ANDERSON *et al.* 2010; GUO *et al.* 2017). The sodium-driven phosphate cotransporters of SLC34 family include three members, NaPi- II a, NaPi- II b and NaPi- II c encoded, respectively, by the gene families SLC34A1, SLC34A2 and SLC34A3 (XU *et al.* 1999). SLC34 family members can mediate transporting divalent inorganic phosphate (HPO4^2-^) into epithelial cells via two or three sodium ions cotransport (WAGNER *et al.* 2014). Type II b sodium phosphate co-transporter (NaPi- II b, gene SLC34A2 localized at chromosome 4p15), is a tissue-specific transporter with molecular weight in 76–110 kDa range expressed in a number of mammalian tissues involved in the metabolism of inorganic phosphorus (ZHENG *et al.* 2009). It has been shown that aberrant expression of NaPi- II b in various malignancies such as gastric cancer (ZHANG *et al.* 2018), lung adenocarcinoma (ZHAO *et al.* 2017), bladder cancer (YE *et al.* 2017), breast cancer (LV *et al.* 2017), hepatocellular carcinoma (LI *et al.* 2016) and ovarian cancer (LIN *et* al. 2015). In ovarian cancer, NaPi- II b was overexpressed comparison to normal tissues and other types of cancer (RANGEL *et al.* 2003). Antibody-drug conjugates targeting SLC34A2 had promising therapeutic effects to OC xenograft models with tolerated toxicity (LIN *et al.* 2015). However, the expression and prognostic effect of NaPi- II b in ovarian cancer remain to be investigated. In the present study, we for the first time extended the research to analyze the expression and prognosis of NaPi- II b in OC based on large databases. We firstly demonstrated that NaPi- II b was significantly overexpressed in OC tissues and predict better survival in OC patients. Therefore, NaPi- II b might be a candidate diagnostic and prognostic biomarker for OC patients.

## MATERIALS AND METHODS

### Oncomine analysis

To determine the expression pattern of SLC34A2 in OC, the datasets in Oncomine database, a powerful platform for data mining, brings cancer microarray data and analysis capabilities to stimulate new discovery were used. SLC34A2 gene was queried and visualized in the database and the results were filtered by selecting OC and OC vs. Normal Analysis. Gene rank threshold was selected as 2 fold change, p-value = 0.001 and top 10%.

### CCLE analysis

Cancer Cell Line Encyclopedia database provides public access analysis and visualization of chromosomal copy number, mRNA expression and massively parallel sequencing data for about1062 human cancer cell lines. The relative mRNA level of SLC34A2 in a various of OC cancers were analyzed by CCLE database.

### The Kaplan-Meier plotter

The prognostic significance of the mRNA expression of SLC34A2 in OC was evaluated using the Kaplan-Meier plotter (www.kmplot.com), an online database can be used to assess the effect of the gene on cancer prognosis.

In this database, data of lung cancer, OC, gastric cancer and breast cancer are available (GYORFFY *et al.* 2010; GYŐRFFY *et al.* 2012; GYORFFY *et al.* 2013; SZÁSZ *et* al. 2016). The patient samples were divided into two cohorts according to the median expression of the gene (high vs. low expression). We analyzed the overall survival (OS) of OC patients by using a Kaplan-Meier survival plot. Briefly, the SLC34A2 gene was uploaded into the database respectively to obtain the Kaplan-Meier survival plots, in which the number-at-risk was shown below the main plot. Log rank p-value and hazard ratio (HR) with 95% confidence intervals were calculated and displayed on the webpage.

## RESULTS

### The expression patterns of SLC34A2 gene based on the Oncomine database

By Oncomine analysis, we investigated mRNA levels of SLC34A2 gene in human cancers including hematogenous malignancies and solid tumors. SLC34A2 was distinctly overexpressed in OC tissues (Figure 1).

**Figure 1.**
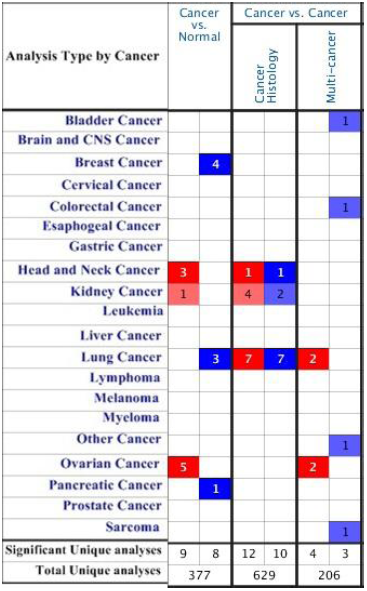
The mRNA levels of SLC34A2 in different tumor types by Oncomine analysis. Red cell represents higher expression and blue cell represents lower expression. The number in each cell represents the number of analyses that meet the threshold of “Gene: SLC34A2; Analysis Type: Cancer vs. Normal Analysis; Cancer Type: ovarian cancer; Data Type: mRNA”.

The relative expression of SLC34A2 was elevated in OC tissues (Figure 2). The respective transcripts of SLC34A2 was 3.604 fold elevated in OC than in normal tissues (Figure 2).

**Figure 2.**
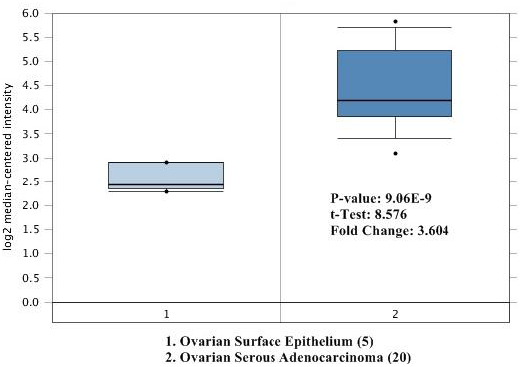
Box plot validating the SLC34A2 expression in normal vs. OC tissue in dataset of Hendrix’s.

### SLC34A2 was distinctively high expressed in OC cell lines from CCLE analysis

The expression of SLC34A2 in OC cell lines respective ranked the top one out of 37 distinct cancer types which represent 1062 cell lines in the *CCLE* databases (Figure 3). SLC34A2 was consistent with that distinctively up-regulated in OC from Oncomine database.

**Figure 3.**
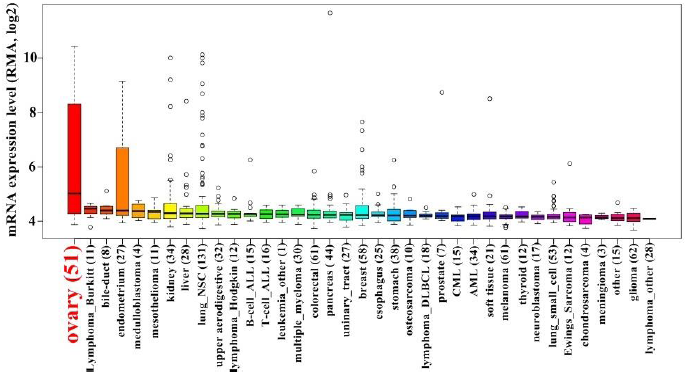
The expressed in cancer cell lines of SLC34A2 from CCLE analysis. The mRNA expression level of SLC39A4 in the GC cell lines vs. normal cell lines ranked the highest among a variety of cancer cell lines (shown in red).

### Association of SLC34A2 expression with survival time for OC patients

To identify the relationship between the patient survival time and mRNA levels of SLC34A2, we analyzed the gene using a Kaplan-Meier plotter. Survival curves were plotted for all OC patients (Figure 4). OC patients with high expression of SLC34A2 showed to be correlated to significantly shorter OS (Overall Survival) (HR=1.36, P=2.1E-5) (Figure 4). These results indicated that the high mRNA expression of SLC34A2 was significantly associated with worse OS in all OC patients.

**Figure 4.**
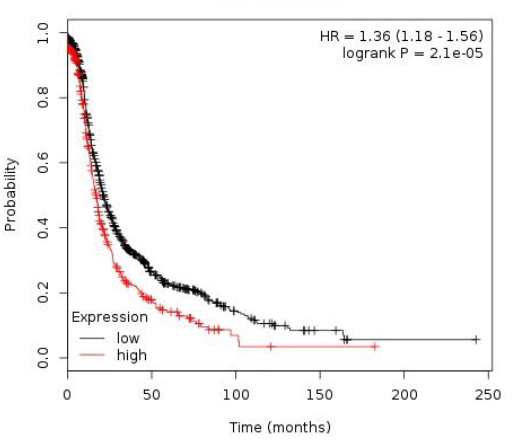
The prognostic values of SLC34A2 expression in OC. High mRNA level of SLC34A2 was associated with worse OS in OC patients.

## DISCUSSION

SLC34A2, widely expressed in human organs, such as lung, liver and small, intestine, ovary, thyroid, breast, and kidney is responsible for maintenance of phosphate homeostasis (XU *et al.* 1999; SABBAGH *et al.* 2009; WAGNER *et al.* 2014). Commonly, dysregulation of SLC34A2 exerts impacts on cellular functions of the initiation and development of human solid malignancies (LI *et al.* 2016; YE *et al.* 2017; ZHANG *et al.* 2017; LIU *et al.* 2018; ZHANG *et al.* 2018). Recent findings have demonstrated that the overexpression of SLC34A2 in maintenance of stem cell–like cells in lung cancer, gastric cancer, and breast cancer through different signaling pathways (MASHIMA *et al.* 2001; TREUTLEIN *et al.* 2014; ZHANG *et al.* 2018). For example, NaPi-IIb affected the progression in gastric cancer stem cell-like cells by regulating miR-25-Gsk3β signaling pathway (ZHANG *et al.* 2018). Li et al. (LI *et al.* 2016) discovered NaPi-IIb activated PI3K/AKT signaling pathway to promote the growth and invasion of hepatocellular carcinoma cells. In bladder cancer, NaPi-IIb drives the malignant growth and progression of cancer cells through upregulating c-Myc (WEN *et al.* 2017). However, the biological function and the underlying molecular mechanisms by which NaPi-IIb regulates features of OC are unknown. It is worth that a monoclonal antibody targeting NaPi-IIb has been developed to treat OC (LIN *et al.* 2015). Meanwhile, a mouse monoclonal antibody called as MX35, is helpful to histological diagnosis and targeted therapy selection (YIN *et al.* 2008).

OC is a malignant neoplasm with high mortality owing to its aggressive nature. Surgical resection increases the 5-year survival rates to 95% in OC patients who are in early stages (FERRO *et al.* 2014). The poor OC prognosis is mainly due to the usually indistinct symptoms, lack of reliable screening test, and tumor resistance to chemotherapy. Thus, explore novel avenues of new tumor markers for OC diagnosis, prognostic, and therapeutic is urgently required.

Here, our study first demonstrated that NaPi-IIb was distinctly high-expressed and predicted worse survival in OC compared with normal controls. Through Oncomine databases and CCLE analysis, the results demonstrated that the relative expression of SLC34A2 elevated significantly in OC vs. normal tissues. Survival analysis suggested that over-expression of SLC34A2 was associated with worse survival in patients with OC. All in all, these results showed the critical role of NaPi-IIb in OC initiation or progression. However, the function and mechanism of NaPi-IIb contribution to the occurrence, development and prognosis of OC still need to be confirmed.

In summary, NaPi-IIb seem to act as promising potential diagnostic and prognostic biomarker in OC. The mRNA expression level of SLC34A2 in OC is distinctly higher than in normal tissues. High expression of SLC34A2 is correlated to worse OS in OC patients. High expression of SLC34A2 might correlate to the occurrence, development and prognosis in OC. However, these results are the bioinformatics analyses, further experimental evidence would be benefit for disclosing NaPi-IIb in the molecular biology of OC.

